# Is degree of sociality associated with reproductive senescence? A comparative analysis across birds and mammals

**DOI:** 10.1101/2020.10.29.360636

**Authors:** Csongor I. Vágási, Orsolya Vincze, Jean-François Lemaître, Péter L. Pap, Victor Ronget, Jean-Michel Gaillard

**Affiliations:** Evolutionary Ecology Group, Hungarian Department of Biology and Ecology, Babe□-Bolyai University, Cluj-Napoca, Romania; Department of Tisza Research, MTA Centre for Ecological Research-DRI, Debrecen, Hungary; CREEC, UMR IRD 224-CNRS 5290-Université de Montpellier, Montpellier, France and CREES Centre for Research on the Ecology and Evolution of Disease, Montpellier, France; Laboratoire de Biométrie et Biologie Évolutive, CNRS, Université Lyon, Villeurbanne, France; Unité Eco-anthropologie (EA), Muséum National d’Histoire Naturelle, CNRS, Université Paris Diderot, Paris, France

**Keywords:** brain size, coloniality, cooperative breeding, life history, reproductive ageing, vertebrates

## Abstract

Our understanding on how widespread reproductive senescence is in the w ild and how the onset and rate of reproductive senescence vary among species in relation to life histories and lifestyles is currently limited. More specifically, whether the species-specific degree of sociality is linked to the occurrence, onset and rate of reproductive senescence remains unknown. Here, we investigate these questions using phylogenetic comparative analyses across 36 bird and 101 mammal species encompassing a wide array of life histories, lifestyles and social traits. We found that female reproductive senescence (1) is widespread and occurs with similar frequency (about two thirds) in birds and mammals; (2) occurs later in life and is slower in birds than in similar-sized mammals; (3) occurs later in life and is slower with an increasingly slower pace of life in both vertebrate classes; and (4) is only weakly associated, if any, with the degree of sociality in both classes after accounting for the effect of body size and pace of life. However, when removing the effect of species differences in pace of life, a higher degree of sociality was associated with later and weaker reproductive senescence in females, which suggests that degree of sociality is either indirectly related to reproductive senescence via the pace of life or simply a direct outcome of the pace of life.

**Subject Areas:** ecology, evolution

## 1. Introduction

Reproductive senescence (or reproductive ageing) – the decline in reproductive performance with increasing age – is widespread in nature [1,2], except for species with indeterminate growth that gain mass and thereby increase fecundity with age [3]. Recent studies have revealed that both the timing and the strength of reproductive senescence is highly variable across species [4,5], although our knowledge is still very limited about how ecological factors and species-specific life history shape variation in either the onset or the rate of reproductive senescence [1,6]. Among these factors, the possible role played by the species-specific degree of sociality has never been investigated.

Sociality is evolutionarily associated with a complex set of life-history traits. Most notably, social species might have longer lifespan and decreased actuarial senescence (see [7–9] for reviews). Indeed, social life in cooperative breeders and colonial species can buffer environmentally-driven mortality risks and might ultimately slow down actuarial senescence (e.g. [10] for a case study on cooperatively breeding Seychelles warblers, *Acrocephalus sechellensis*), even if the relationship between sociality and actuarial senescence is likely to be complex and might differ both within and among species [7]. However, the association between social life and the occurrence, onset and rate of reproductive senescence has never been investigated so far, although similar relationship with the intensity of senescence is expected for survival and reproduction. We aimed here to fill this knowledge gap using the most comprehensive comparative analyses performed to date across bird and mammal species.

Within populations, there is a large variation among individuals in their sociability. Even within highly social species, some individuals are more connected to others, while some have few and loose social interactions with conspecifics (e.g. variation according to social status and environmental context in spotted hyena, *Crocuta crocuta* [11]; variation with age in yellow-bellied marmots, *Marmota flaviventer* [12]; variation in early social development in bottlenose dolphin, *Tursiops* sp. [13]). In cooperative breeders, most of the individuals are social during at least part of their life [14]. Nevertheless, even within these populations, individuals are not equally social and they differ in the amount of help they receive and provide. The evolutionary hypotheses explaining why social individuals should display a weaker senescence than solitary ones [7] are rooted in the principle of allocation [15]. This principle states that increased allocation of finite resources to a given biological function (e.g. reproduction) compromises allocation to a competing function (e.g.somatic maintenance that promotes survival) [16]. Increased allocation of resources to reproduction early in life, which is favoured by natural selection in growing populations [17], is expected to have detrimental consequences in terms of actuarial and/or reproductive senescence [6]. This trade-off is predicted by both antagonistic pleiotropy and disposable soma theories of ageing [18,19], and is well supported by current empirical evidence [20,21]. For instance, male red deer (*Cervus elaphus*)allocating substantial resources to sexual competition during early life show a steeper rate of reproductive senescence in late life ([22]; see also [23] for examples in birds). How social lifestyle may buffer against such costs? For instance, helpers in cooperative breeders reduce the workload of reproducers according to the load-lightening hypothesis [24]. Thus, the principle of allocation, a key concept of life-history evolution [16,25], explains how senescence can either increase due to adelayed cost of high performance during early life [20,26] or decrease thanks to a reduced reproductive effort required under high degree of sociality (e.g. the presence of helpers [24]).Assuming that these processes can explain the variance in senescence observed at the inter-specific level, two main hypotheses can be proposed to expect a negative covariation between the degree of sociality and reproductive senescence:

> (H1) Given the inevitable costs of reproduction [27,28], a high reproductive effort observed in a given species should lead to an earlier and/or faster reproductive senescence [20]. For a given reproductive effort, a higher degree of sociality in a given species might facilitate there productive duties of individuals and therefore reduce directly the costs of reproduction [7]and ultimately shape the senescence patterns of that species [7,21]. Thus, the mitigation of reproductive cost by a social mode of life should lead to postponed onset and/or decelerated rate of reproductive senescence of a given species

> (H2) The degree of sociality can drive the evolution of reproductive senescence in a given species indirectly through decreasing adult mortality risk, thereby slowing down the pace of life. Life-history theory postulates that a decreased rate of environmentally-driven mortality should favour slower growth rate, longer time to maturation, older age at first reproduction and reduced allocation to reproduction by young adults [16], as well as later onsets and slower rates of both actuarial and reproductive senescence [6,29]. Indeed, sociality has been shown to mitigate multiple forms of environmentally driven mortality risks (e.g. starvation, predation). Thus, the presence of social partners in a given species is associated with a slowing down of the pace of life, which leads to delayed and decelerated reproductive senescence in both mammals and birds [5].

Under both hypotheses, reproductive senescence should be less pronounced in species with a higher degree of sociality by involving either a direct response to reproductive effort at each reproductive attempt (H1) or indirectly through a slower pace of life selecting for a lower reproductive effort early in life (H2). If the degree of sociality is directly associated with reproductive senescence (H1), we predict a substantial effect of the degree of sociality even after the effects of allometry and pace of life on reproductive senescence are accounted for. If the degree of sociality is indirectly associated with reproductive senescence via the pace of life (H2), we predict no detectable effect of the degree of sociality once the effects of allometry and pace of life are accounted for.

Here, we modelled age-specific changes in reproductive traits at the species level and tested whether the degree of sociality accounts for the variation in the occurrence, onset and rate of reproductive senescence observed across birds and mammals (*n* = 36 and 101 species, respectively). The age when reproductive performance starts to decline marks the onset, while the slope of the age-specific decline in reproductive performance fitted from the onset expresses the rate. We followed strict statistical rules to assess whether reproductive senescence occurred (see Methods) and estimated onset and rate only for species in which it did occur (i.e. species with a statistically significant decrease of reproductive performance with increasing age). We accounted for the confounding effect of phylogenetic inertia, allometric constraints and species’ ranking on the slow– fast continuum of life histories (i.e. pace of life) in our phylogenetic comparative analyses, as all these processes are known to shape variation in senescence [5].

## 2. Methods

### (a) Female reproductive senescence data

As age-specific reproductive output is easier to measure in females than in males (e.g. due to extra-pair offspring often produced by males; [30]) and has been reported in a much higher number of vertebrate species, we focus on the reproductive ageing of females in both birds and mammals. Reproductive senescence parameters of 101 wild or semi-captive mammal species were taken from [31]. This data set includes the presence/absence of reproductive senescence and, for species with evidence of senescence, the age at onset and the rate of reproductive senescence. All those parameters were estimated from age-specific birth rates (i.e. number of female offspring alive at birth that are produced by a female of age *x*, tabulated as *m*_*x*_ in a life table) extracted from published life tables or graphs using WebPlotDigitizer (https://automeris.io/WebPlotDigitizer/). The acquisition of age-specific reproductive data for mammals is fully detailed in [31]. In cooperatively breeding mammals, age-specific reproductive data were collected for dominant females (e.g. [32]), as subordinate females generally have no access to reproduction.

In birds, we conducted a systematic literature search of age-specific changes in reproductive traits in wild populations to extract data similar to those obtained for mammals (see Electronic Supplementary Material, ESM for search methods). Unlike in mammals, age-specific birth rates (i.e. the *m*_*x*_ parameter) were seldom reported in bird studies because the probability of breeding – necessary for birth rate calculations – is often unknown. Therefore, to increase the number of species, we also included studies that reported age-specific number of hatchlings or number of fledglings per female when birth rates could not be extracted or computed. Some studies reported standardized values (i.e. normalized values or residuals from models) instead of raw values of age-specific reproduction. We included those studies in our analyses and controlled for the effect of analysing standardized data (yes/no). When reproductive data were reported for multiple populations of the same species, we only included the study with the largest sample size, as done in mammals [31]. To estimate reproductive senescence parameters, we accounted for differences in the age-specific sample sizes, as done in mammals [31]. We used the original age-specific sample size when reported in the original studies, and we calculated the number of females expected to be alive at age *x* from the observed age distribution of females when sample sizes were not reported. We collected female reproductive data for 36 avian species (see ESM ‘Data set’).

Age-dependent reproductive traits in birds followed similar distributions to mammalian ones. Hence, we computed reproductive senescence parameters in birds using the same methods as in mammals. Briefly (see [31] for further details), four different age-dependent models (i.e. constant model, linear model, threshold model with one threshold and two linear segments, and threshold model with two thresholds and three linear segments) weighted by the age-specific sample size were fitted on the reproductive data using the R package ‘segmented’ [33]. The final model was selected using Akaike’s Information Criterion (AIC) (see the Methods in the ESM for model selection procedure and ESM ‘Model selection’ for the AIC values associated to each alternative senescence models; see also ESM ‘Segmented’ for the segmented fits of the selected models plotted separately for each bird species; similar table and plots for mammals can be found in [31]). Based on the selected model, different procedures were used to infer reproductive senescence from the slope of the different linear segments and their associated standard error. When reproductive senescence occurred (i.e. slope of one of the segments < 0), the rate and the onset of reproductive senescence were reported as the slope of the linear segment and the age corresponding to the beginning of the segment, respectively. Using this procedure, we detected reproductive senescence for most of the bird species for which it was observed in the original studies from which the data were extracted. Only minor discrepancies were found mostly due to the use of different statistical methods (see ESM ‘Occurrence’ for a comparison of the results found on reproductive senescence using our standardized procedure against the results found in the original studies; a similar comparison for mammals can be found in [31]).

### (b) Life-history traits

To assess the relationship between the degree of sociality and reproductive senescence, we first had to account for inter-specific differences in body size and biological time [34], which structure most life-history variation across vertebrates [35]. Body mass is a reliable measure of species-specific size that shapes age-specific reproductive and survival rates via allometric effects. Thus, small bird and mammal species display both earlier and steeper reproductive senescence than large ones [5]. Likewise, for a given size, slow-living species display both later and slower reproductive senescence, an effect well illustrated by the comparison of similar-sized birds and mammals [5]. Generation time is the most appropriate metric to position species on the slow–fast continuum of life histories [36]; however, data to accurately measure generation time were missing for many of the species studied here [37]. Thus, instead of generation time, we used a compound of the age at first reproduction and maximum longevity observed in the focal case study to measure species-specific pace of life (see below). In birds, we collected data on female body mass from [38], age at first reproduction and longevity from the same papers including age-specific reproduction data (ESM ‘Data set’), while in mammals data of the same traits were retrieved from [31].

### (c) Sociality traits

The social environment varies considerably across species and this diversity can have vast evolutionary consequences [39]. We use four simple sociality traits (i.e. coloniality, parental cooperation, cooperative breeding and relative brain size; see also [40,41]) to assess the species-specific degree of sociality (table 1) and test whether these traits are associated or not with the occurrence, rate and onset of reproductive senescence across birds and mammals. These four proxies of sociality cover different ranges of degree of sociality. For instance, cooperative breeders often live in social systems with more complex social interactions than colonial ones, and therefore imply different costs and benefits to the individuals. The diversity of social traits we use in this study makes possible to assess whether social lifestyle in general or specific social systems in particular are associated with reproductive senescence, if any.

**Table 1.** Sociality traits and their meaning in terms of degree of sociality.

| sociality trait | degree of sociality |  |
| --- | --- | --- |
|  | low | high |
| colonial breeding (birds) | no | yes |
| parental cooperation (birds) | female care | female & male care |
| cooperative breeding (mammals) | no | yes |
| relative brain size (birds and mammals) | small | large |

We used three sociality traits in birds (i.e. presence/absence of coloniality, parental cooperation and relative brain size). Both the degree of sociality and the use of social information are higher in species breeding in large and dense colonies of non-kin individuals as compared with solitarily breeding ones [42]. Coloniality has been considered as a proxy of sociality degree in studies of longevity across bird species [40]. We used parental cooperation as a metric of the degree of sociality, which reflects whether female-only, male-only or shared female–male parental care is typical for a given bird species [43]. Family, where individuals form short-term pair bonds during breeding and raise their offspring cooperatively (i.e. have biparental care) is the simplest social system, and species with biparental care display a higher degree of sociality than species in which only females care for their young [41]. This metric is relevant in birds because it influences the reproductive costs of females and thus is likely to modulate female reproductive senescence parameters. Biparental care is the most common form of social behaviour between unrelated individuals in birds, with over 90% of all living birds being biparental [44]. The presence of cooperative breeding was not considered in birds due to the low number of species with regular cooperative breeding in our data set.

We used two sociality traits in mammals (i.e. presence/absence of cooperative breeding and relative brain size). The degree of sociality is considered high (i.e. implying frequent and complex social interactions among individuals) in species living in small cooperative breeding groups with helpers as compared with non-cooperatively breeding ones. Cooperative breeders have the most intense social system among mammals [45]. Because coloniality cannot be defined with confidence in mammals, but in a few species only (e.g. in black-tailed prairie dogs, *Cynomys ludovicianus* [46]), we had to omit this sociality trait in this vertebrate class.

The relative brain size (i.e. brain size for a given body size) was used for both birds and mammals. Relative brain size is higher in species with high degree of social bonding (e.g. primates and whales/dolphins) or reproductive pair bonding (e.g. monogamous carnivores and ungulates, bats and birds) [41], making possible its use to measure the degree of sociality [41,47]. Quantifying the degree of sociality in comparative studies encompassing species with a large range of life-history strategies is far from trivial, which leads most comparative studies to use only proxies of sociality instead of accurate metrics.

In birds, we collected data on brain mass from [48], presence/absence of coloniality from [49] and parental cooperation during breeding from [43] (see ESM ‘Data set’). In the latter source, parental cooperation was separately quantified for the pre- and post-hatching periods, which are highly correlated (Pearson correlation *r* = 0.76, df = 29, *t* = 6.24, *p* < 0.0001). We calculated the average of these two periods (henceforth parental cooperation) reflecting the sex bias in parental care during breeding. Values range from –1 (exclusive female care) to 1 (exclusive male care), with 0 reflecting an equal share of parental duties between sexes. Coloniality had a perfect overlap with marine environment in our bird data set, as all colonial species are seabirds and all solitary ones are terrestrial, which reflects a strong phylogenetic bias and a limitation of our coloniality data (see Discussion). In mammals, data on brain mass were obtained from [50], presence/absence of cooperative breeding from [51] and we completed species with lacking information with additional sources (see ESM ‘Data set’).

### (d) Statistical analyses

All analyses were performed in R version 4.0.1 [52]. To make meaningful inferences about the effect of body size, pace of life and degree of sociality on reproductive senescence, all models were controlled for phylogenetic inertia. In birds we used a rooted, ultrametric consensus tree built using the SumTrees Python library [53] based on 1,000 trees. These trees were obtained from birdtree.org [54] using the Hackett backbone tree [55]. For mammals, we used a published phylogenetic super-tree (see also [56]).

Female body mass, age at first reproduction, longevity and brain mass were highly correlated across both bird and mammal species (table S1). Therefore, to avoid multicollinearity problems, we conducted a phylogenetically-controlled principal component analysis (PPCA) as implemented in R package ‘phytools’ [57] on the first three traits (all log-transformed) separately for birds and mammals. We retained the first two phylogenetic principal components (PPCs), where the first PPC is a size component (hereafter PPC size), which explained 69% and 79% of variation in birds and mammals, respectively, and the second PPC is a pace of life component (hereafter PPC pace), which explained additional 23% and 12% of variation in birds and mammals, respectively (table S2). Larger values indicate larger body mass (PPC size) and slower pace of life (PPC pace), respectively (table S2). PPC size and PPC pace were used in the subsequent analyses to control for allometry and pace of life, respectively. Given that we were specifically interested in the effect of relative brain size (a proxy measure of the degree of sociality) on reproductive senescence, we did not include brain size in the PPCA. Nonetheless, to avoid collinearity of brain size with PPC size, we estimated relative brain size as residuals of a standard major axis regression (as implemented in R package ‘lmodel2’) between log-transformed brain size and PPC size and used this measure in the multifactorial models.

To explore variation in reproductive senescence patterns, we used phylogenetic logistic regressions for evidence of reproductive senescence and phylogenetic linear regressions separately for onset and rate of reproductive senescence as implemented in R package ‘phylolm’ [58]. Age at onset and the absolute value of the rate of reproductive senescence were log-transformed prior to the analysis. In birds, for each senescence metric, the reproductive trait used to assess reproductive senescence (i.e. birth rate *m*_*x*_, number of hatchlings or number of fledglings) and the presence/absence of coloniality were tested as fixed factors, while PPC size, PPC pace, residual brain size and parental cooperation were included as covariates. We did not need to account for either the hunting status (because no bird species in the data set is hunted) or the data quality (because all bird studies were based on longitudinal data and only included known-aged individuals). Similarly to analysis in mammals (see [31]), we tested whether the probability to detect reproductive senescence in birds was influenced by the sample size (i.e. total number of reproductive records in the population; log-transformed in the analysis; ESM ‘Data set’). For the rate of reproductive senescence, the effect of data standardization (yes/no) was also tested. In mammals, for each senescence metric, data quality (transversal/longitudinal), hunting status (hunted/not hunted) and presence/absence of cooperative breeding were included as fixed factors, while PPC size, PPC pace and residual brain size were included as covariates. In both birds and mammals, the effect of age at onset of reproductive senescence (log-transformed) was also tested in models of rate of reproductive senescence because a negative correlation is expected to occur [31]. Non-linear effects of PPC size and PPC pace were also modelled in both bird and mammal models using second-degree orthogonal polynomials, but were only retained in the model when their inclusion decreased AIC values by > 2 compared with the initial model without the polynomials. In no case where a quadratic model was selected over the linear model did a cubic model outperform the quadratic model, meaning that a second-order polynomial satisfactorily accounted for observed non-linear relationships. Sample size varied across models because some variables (e.g. brain size) had missing values in certain species and rate as well as onset of senescence were only analysed for species in which evidence of reproductive senescence was detected. To test H2 according to which the effect of the degree of sociality acts indirectly through slowing down the pace of life, we reran all the above-mentioned analyses after removing PPC pace from the models.

Due to the limited number of bird species, we adopted an AIC-based stepwise forward model selection procedure to avoid over-parametrization of models. As a first step, an intercept model was constructed for each dependent variable. In the second step, each explanatory variable (except metrics of sociality) was added one by one to this model and the model with the smallest AIC value (if ΔAIC < 2) was further elaborated until adding extra variables did not decrease AIC value by > 2. This model is referred as the base model. If any of the single-predictor models had ΔAIC < 2, the intercept model was considered as the base model. In the third step, to test the association between the degree of sociality and reproductive senescence, the sociality traits were added one by one to the base model and the change in AIC was checked (table S3). Given that relative brain size and parental cooperation had missing values for some species, when testing their effect on reproductive senescence metrics, their corresponding base models were refitted for the subset of species with the full set of available data. These models are presented in table S3, while table 2 shows the ANOVA results of the base models presented in table S3.

**Table 2.** Base models of occurrence (*a*), rate (*b*) and onset (*c*) of reproductive senescence in birds (see table S3 for AIC-based stepwise forward model selection in birds). PPC size and PPC pace are the phylogenetic principal components describing size and pace of life, respectively. Models on the left include pace of life, while those on the right do not include pace of life. The statistically significant linear or polynomial effect of pace of life (PPC pace and poly(PPC pace), respectively) is marked in bold in models on the left side. Social traits are italicized and those with statistically significant effect are italicized and marked in bold. α and λ – phylogenetic signal; AIC – Akaike Information Criterion; *n* – sample size (number of species).

| including pace of life (PPC pace) |  |  |  | excluding pace of life (PPC pace) |  |  |  |
| --- | --- | --- | --- | --- | --- | --- | --- |
| (a) occurrence of reproductive senescence |  |  |  | (a) occurrence of reproductive senescence |  |  |  |
| predictors | $\beta$ (s.e.) | $z$ | $p$ | predictors | $\beta$ (s.e.) | $z$ | $p$ |
| intercept | 0.57 (0.37) | 1.53 | 0.1254 | intercept | 0.57 (0.37) | 1.53 | 0.1254 |
| model stats: $\alpha = 0.1860$, AIC = 49.04, $n = 36$ | | | | model stats: $\alpha = 0.1860$, AIC = 49.04, $n = 36$ | | | |
| (b) log rate of reproductive senescence |  |  |  | (b) log rate of reproductive senescence |  |  |  |
| predictors | $\beta$ (s.e.) | $t$ | $p$ | predictors | $\beta$ (s.e.) | $t$ | $p$ |
| intercept | -3.37 (0.38) | 8.89 | < 0.0001 | intercept | -3.14 (0.45) | 6.98 | < 0.0001 |
| poly(PPC size, 2)1 | -3.23 (0.81) | 4.00 | 0.001 | poly(PPC size, 2)1 | -1.10 (0.88) | 1.25 | 0.2291 |
| poly(PPC size, 2)2 | -1.41 (0.69) | 2.04 | 0.0584 | poly(PPC size, 2)2 | -2.25 (0.75) | 2.98 | 0.0088 |
| **PPC pace** | **-0.89 (0.23)** | **3.79** | **0.0016** | repr. trait (no. hatchlings) | 1.40 (0.58) | 2.40 | 0.0288 |
| repr. trait (no. hatchlings) | 0.97 (0.54) | 1.80 | 0.091 | repr. trait (no. fledglings) | 1.08 (0.45) | 2.38 | 0.0299 |
| repr. trait (no. fledglings) | 1.12 (0.41) | 2.73 | 0.0149 | **coloniality** | **-1.06 (0.35)** | **3.01** | **0.0084** |
| model stats: $\lambda = 0.000$, AIC = 53.71, $n = 22$ | | | | model stats: $\lambda = 0.000$, AIC = 57.93, $n = 22$ | | | |
| (c) log onset of reproductive senescence |  |  |  | (c) log onset of reproductive senescence |  |  |  |
| predictors | $\beta$ (s.e.) | $t$ | $p$ | predictors | $\beta$ (s.e.) | $t$ | $p$ |
| intercept | 2.16 (0.10) | 22.65 | < 0.0001 | intercept | 2.05 (0.14) | 14.16 | < 0.0001 |
| PPC size | 0.32 (0.04) | 7.37 | < 0.0001 | PPC size | 0.24 (0.05) | 4.88 | 0.0001 |
| **PPC pace** | **0.39 (0.14)** | **2.88** | **0.0096** | **coloniality** | **0.41 (0.21)** | **1.96** | **0.0648** |
| model stats: $\lambda = 0.000$, AIC = 30.19, $n = 22$ | | | | model stats: $\lambda = 0.000$, AIC = 34.10, $n = 22$ | | | |

Given the large sample size in mammals, we present the full models with all explanatory variables entered simultaneously (table 3). Consequently, the final sample size is 88 mammalian species (out of 101 species) because brain size data were missing for 13 species. However, repeating the analyses by excluding brain size and keeping only cooperative breeding as sociality trait, which is available for the entire species pool, the results of cooperative breeding remain unchanged (results not shown

**Table 3.** Full models of occurrence (*a*), rate (*b*) and onset (*c*) of reproductive senescence in mammals. PPC size and PPC pace are the phylogenetic principal components describing size and pace of life, respectively. Models on the left include pace of life, while those on the right do not include pace of life. The statistically significant linear or polynomial effect of pace of life (PPC pace and poly(PPC pace), respectively) is marked in bold in models on the left side. Social traits are italicized and those with statistically significant effect are italicized and marked in bold. α and λ – phylogenetic signal; AIC – Akaike Information Criterion; *n* – sample size (number of species).

| including pace of life (PPC pace) |  |  |  | excluding pace of life (PPC pace) |  |  |  |
| --- | --- | --- | --- | --- | --- | --- | --- |
| (a) occurrence of reproductive senescence |  |  |  | (a) occurrence of reproductive senescence |  |  |  |
| predictors | $\beta$ (s.e.) | $z$ | $p$ | predictors | $\beta$ (s.e.) | $z$ | $p$ |
| intercept | 1 (0.44) | 2.26 | 0.0236 | intercept | 1.13 (0.43) | 2.62 | 0.0088 |
| quality (transversal) | -1.29 (0.51) | 2.51 | 0.0119 | quality (transversal) | -1.29 (0.52) | 2.48 | 0.0132 |
| hunted (yes) | -0.31 (0.6) | 0.51 | 0.6074 | hunted (yes) | -0.4 (0.6) | 0.67 | 0.5058 |
| PPC size | 0.23 (0.1) | 2.18 | 0.0294 | PPC size | 0.23 (0.1) | 2.27 | 0.0233 |
| PPC pace | -0.01 (0.4) | 0.03 | 0.9755 |  |  |  |  |
| *residual brain size* | *1.01 (0.64)* | *1.59* | *0.1129* | *residual brain size* | *0.99 (0.61)* | *1.62* | *0.1062* |
| *cooperative breeding (yes)* | *0.1 (0.82)* | *0.12* | *0.9077* | *cooperative breeding (yes)* | *0.02 (0.83)* | *0.02* | *0.9855* |
| model stats: $\alpha$ = 0.0434, AIC = 109.46, $n$ = 88 | | | | model stats: $\alpha$ = 0.0544, AIC = 107.28, $n$ = 88 | | | |
| (b) log rate of reproductive senescence |  |  |  | (b) log rate of reproductive senescence |  |  |  |
| predictors | $\beta$ (s.e.) | $t$ | $p$ | predictors | $\beta$ (s.e.) | $t$ | $p$ |
| intercept | -2.12 (0.59) | 3.58 | 0.0008 | intercept | -1.6 (0.54) | 2.97 | 0.0045 |
| log onset of senescence | -0.03 (0.27) | 0.12 | 0.9083 | log onset of senescence | -0.33 (0.24) | 1.37 | 0.1758 |
| quality (transversal) | -0.06 (0.28) | 0.21 | 0.8370 | quality (transversal) | -0.07 (0.3) | 0.23 | 0.8170 |
| hunted (yes) | 0.54 (0.35) | 1.56 | 0.1260 | hunted (yes) | 0.56 (0.36) | 1.57 | 0.1220 |
| PPC size | -0.43 (0.06) | 6.73 | < 0.0001 | PPC size | -0.44 (0.08) | 5.37 | < 0.0001 |
| **poly(PPC pace, 2)1** | **-4.45 (1.35)** | **3.28** | **0.0019** |  |  |  |  |
| **poly(PPC pace, 2)2** | **-2.41 (1.08)** | **2.24** | **0.0295** |  |  |  |  |
| *residual brain size* | *-0.14 (0.31)* | *0.46* | *0.6483* | *residual brain size* | *-0.64 (0.32)* | *2.03* | *0.0475* |
| *cooperative breeding (yes)* | *1.3 (0.43)* | *3.05* | *0.0037* | *cooperative breeding (yes)* | *0.85 (0.49)* | *1.71* | *0.0927* |
| model stats: $\lambda$ = 0.0417, AIC = 161.09, $n$ = 58 | | | | model stats: $\lambda$ = 0.2794, AIC = 171.72, $n$ = 58 | | | |

| (c) log onset of reproductive senescence |  |  |  | (c) log onset of reproductive senescence |  |  |  |
| --- | --- | --- | --- | --- | --- | --- | --- |
| predictors | $\beta$ (s.e.) | $t$ | $p$ | predictors | $\beta$ (s.e.) | $t$ | $p$ |
| intercept | 1.81 (0.18) | 10.33 | < 0.0001 | intercept | 1.87 (0.24) | 7.63 | < 0.0001 |
| quality (transversal) | -0.17 (0.13) | 1.3 | 0.1995 | quality (transversal) | -0.17 (0.14) | 1.18 | 0.2444 |
| hunted (yes) | 0.15 (0.16) | 0.94 | 0.3494 | hunted (yes) | 0.04 (0.17) | 0.21 | 0.8311 |
| PPC size | 0.18 (0.04) | 4.8 | < 0.0001 | PPC size | 0.23 (0.05) | 5 | < 0.0001 |
| **PPC pace** | **0.54 (0.13)** | **4.11** | **0.0001** |  |  |  |  |
| *residual brain size* | *0.1 (0.17)* | *0.56* | *0.5811* | ***residual brain size*** | ***0.38 (0.19)*** | ***2.01*** | ***0.0499*** |
| *cooperative breeding (yes)* | *-0.13 (0.25)* | *0.55* | *0.5861* | *cooperative breeding (yes)* | *0.12 (0.26)* | *0.47* | *0.6375* |
| model stats: $\lambda = 0.6053$, AIC = 85.94, $n = 58$ | | | | model stats: $\lambda = 0.8102$, AIC = 98.5, $n = 58$ | | | |

## 3. Results

### (a) Occurrence of senescence in birds and mammals

Reproductive senescence was detected in 61% (22 out of the 36 species) of bird species and 68% (69 out of 101 species) of mammal species. The occurrence of reproductive senescence was similar in birds and mammals (Chi-squared test *χ*^2^ = 0.34, df = 1, *p* = 0.562). The probability of detecting reproductive senescence tended to increase with sample size in birds (*β* ± s.e. = 0.43 ± 0.31, *p* = 0.16), but this effect was not statistically significant (as opposed with mammals, see [31]).

### (b) Allometry, pace of life and the degree of sociality in birds

Results of occurrence, rate and onset of reproductive senescence in birds are presented in table 2 and table S3.

The occurrence of reproductive senescence in birds was unrelated to body size and pace of life, and was independent of the reproductive trait used to assess reproductive senescence. None of the sociality traits was associated with the probability to detect reproductive senescence (table 2*a*, table S3*a*).

The rate of reproductive senescence decreased non-linearly with increasing body size (linear term: *β* ± s.e. = –3.23 ± 0.81; quadratic term: *β* ± s.e. = –1.41 ± 0.69), decreased linearly with an increasingly slower pace of life (*β* ± s.e. = –0.89 ± 0.23), and varied among reproductive traits used to assess reproductive senescence. The rate of reproductive senescence was the slowest when using birth rates, intermediate when using the number of hatchlings and fastest when using the number of fledglings. Data standardization did not explain substantial variation in the rate of reproductive senescence either. The rate of reproductive senescence tended to decrease with increasingly later onset of senescence, although this effect was not statistically significant. None of the sociality traits was associated with the rate of reproductive senescence, which does not support H1. Once the marked effect of pace of life was removed from the model, the rate of reproductive senescence was slower in colonial birds than in solitary breeders (*β* ± s.e. = –1.06 ± 0.35), in support of H2 (table 2*b*, table S3*b*).

The onset of reproductive senescence increased linearly with both body size (*β* ± s.e. = 0.32 ± 0.04) and slower pace of life (*β* ± s.e. = 0.39 ± 0.14). None of the sociality traits was related to the age at onset of reproductive senescence, which does not support H1. Once the strong effect of pace of life was removed from the model, the onset of reproductive senescence was later in colonial birds than in solitary species (*β* ± s.e. = 0.41 ± 0.21), in support of H2 (table 2*c*, table S3*c*).

These results do not support H1, but do support H2, which involves an indirect relationship between degree of sociality and both the rate and onset of reproductive senescence via a slowing down of the overall pace of life in species with higher degree of sociality.

### (C) Allometry, pace of life and the degree of sociality in mammals

Results of occurrence, rate and onset of reproductive senescence in mammals are presented in table 3.

Reproductive senescence was more likely to be detected when data originated from longitudinal rather than transversal studies (*β* ± s.e. = –1.29 ± 0.52). Larger-sized mammals were more likely to experience reproductive senescence than smaller ones (*β* ± s.e. = 0.23 ± 0.10). Neither relative brain size, nor cooperative breeding was related to the probability to detect reproductive senescence in mammals (table 3*a*).

The rate of reproductive senescence decreased linearly with increasing body size (*β* ± s.e. = –0.43 ± 0.06) and non-linearly with increasingly slower pace of life (linear term: *β* ± s.e. = –4.45 ± 1.35; quadratic term: *β* ± s.e. = –2.41 ± 1.08). Contrary to H1, cooperative breeding mammals had higher rates of reproductive senescence as compared with non-cooperative species (*β* ± s.e. = 1.3 ± 0.43; figure 1), while relative brain size was unrelated to the rate of reproductive senescence. When the marked effect of pace of life was removed from the model, species with larger relative brain size had slower rate of reproductive senescence (*β* ± s.e. = –0.64 ± 0.32), in support to H2. When the pace of life was not controlled for, however, the relationship between cooperative breeding and the rate of reproductive senescence disappeared, which does not support H1 (table 3*b*).

**Figure 1.**
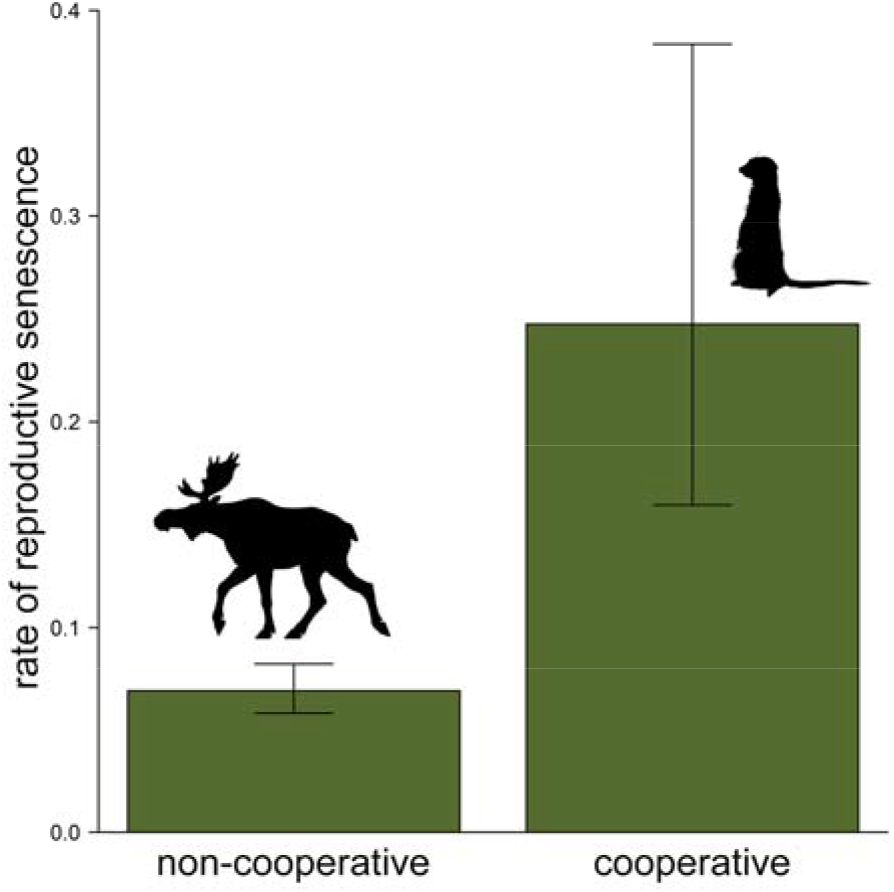
Difference in the rate of reproductive senescence (± s.e.) between cooperative breeding and non-cooperative breeding mammals. Estimated marginal means are plotted, which were extracted from the full model of rate of reproductive senescence with pace of life included among the predictors (see table 3*b*).

The age at onset of reproductive senescence increased linearly with both body size and increasingly slower pace of life (*β* ± s.e. = 0.23 ± 0.05). Neither relative brain size, nor cooperative breeding was related to the age at onset of reproductive senescence in mammals, which does not support H1. Once the marked effect of pace of life was removed from the models, species with large relative brain size showed a later onset of senescence than species with small relative brain size (*β* ± s.e. = 0.38 ± 0.19; table 3*c*), in support to H2.

As in birds, these results do not support H1, but support H2 that involves an indirect relationship between degree of sociality and the rate and onset of reproductive senescence via a slowing down of the overall pace of life in species with higher degree of sociality.

### (d) Comparing reproductive senescence between birds and mammals

In both classes, the rate of reproductive senescence tended to decrease with increasingly later onset of reproductive senescence, with the same apparent strength (figure 2*a*). However, the relationship was statistically significant only in mammals likely because of a lack of power (smaller sample size) in birds. When looking at the allometric relationships, the rate of reproductive senescence decreased (figure 2*b*) and the onset of reproductive senescence occurred later with increasing size in both birds and mammals (figure 2*c*). Interestingly, for a given body mass, mammals displayed both steeper and earlier reproductive senescence than birds did (figure 2*b*,*c*), which is in line with the common view that birds senesce less than similar-sized mammals.

**Figure 2.**
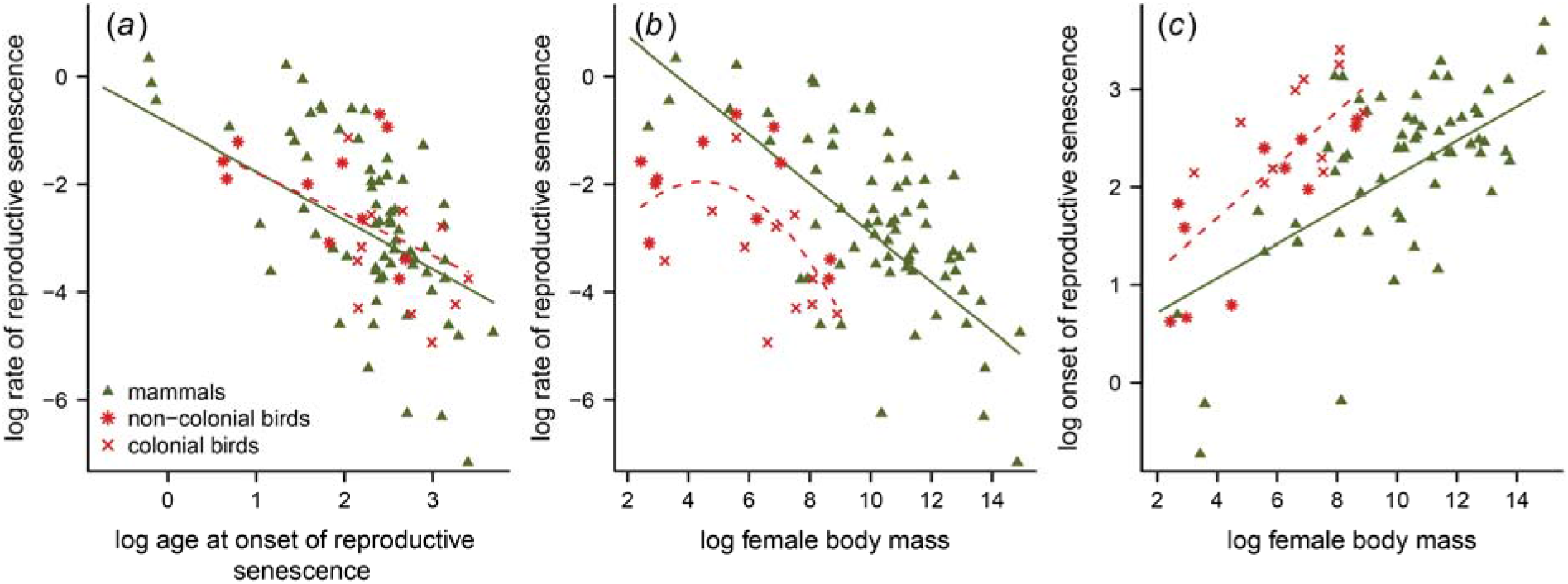
Association between (*a*) age at onset and rate of reproductive senescence, (*b*) female body mass and rate of reproductive senescence, and (*c*) female body mass and age at onset of reproductive senescence across bird species (* non-colonial, × colonial) and mammal species (▲). Female body mass was used here to measure body size because it captures the differences in size range between birds and mammals unlike PPC size, and it is very strongly correlated with PPC size (table S2). Slopes were obtained from single-predictor phylogenetic regressions between the plotted variables (dashed line for birds, continuous line for mammals). Polynomial effect of size is plotted only for rate of senescence in birds because the quadratic term was only statistically significant in this model (see tables 2 and 3).

## 4. Discussion

A previously published review revealed an increasing number of case studies reporting reproductive senescence in the wild [2]. Here, we quantified the occurrence of female reproductive senescence on the largest species-level data set so far compiled on birds and mammals. We found that the proportion of species that display detectable reproductive senescence is similar in avian (0.61; present study) and mammalian (0.68; [31]) species. Interestingly, these proportions are similar to those reported in a previous comparative study of 19 species of birds and mammals (0.65; [5]). However, as the current prevalence of reproductive senescence is likely to be under-estimated (see [31] for further discussion), the biological meaning of these values is disputable. Nevertheless, these studies together emphasize that reproductive senescence is the rule rather than the exception, at least in endotherm vertebrates. The positive effect of sample size on the probability of detecting senescence in mammals (see [31]) constitutes a limitation of our analyses, although this limitation is not detectable for birds, likely due to the smaller data set in this class.

Our findings highlight that birds display a later onset and a slower rate of reproductive senescence as compared with similar-sized mammals. Note that the strength of senescence in birds increases with offspring developmental phase considered for senescence estimates (i.e. from birth rate to number of hatchlings and number of fledglings), while for mammals, senescence was only computed for birth rates. Therefore, we expect that the differences in onset and rate of reproductive senescence between the two classes would be even stronger if weaning success were also considered in mammals. This notion is supported by a recent review showing that maternal effect senescence (i.e. an increasing offspring mortality with mother age, termed Lansing effect) is very common in mammals, while birds being conspicuous exceptions [59]. The more intense reproductive senescence in mammals than in same-sized birds we report matches the class differences reported in longevity (i.e. birds live c.a. 1.5 times longer than similar-sized mammals [60]). Birds also display a much slower pace of life, and, for a given pace of life, birds and mammals of a given size have similar senescence patterns [5]. This suggests that the *modus operandi* of senescence has a deep evolutionary root and is mostly shaped by allometric constraints and pace of life. To test whether the differences between the two classes are explained by flight capacity in birds, and hence their lower environmentally driven mortality, a comparison of reproductive senescence between birds and flying mammals would be promising (see [61] for longevity).

In line with previous observations for other biological times (e.g. longevity, gestation length; see [62]), we found strong effects of allometry and pace of life on both the rate and the onset of reproductive senescence in both birds and mammals. The heavier and slower-paced a species is, the more postponed and slower its senescence is. Both senescence metrics correspond to biological times with a dimension of time for the onset of senescence and a dimension of frequency (i.e. inverse of time) (sensu [34]) for the rate of senescence, which explains the negative relationship we found between the rate and the onset of senescence across birds and mammals. Our analyses thus provide a first evidence that these senescence metrics can be interpreted as life-history traits describing the speed of the life cycle of a given species, alike development time [63], age at first reproduction [64] or longevity [65], which have been much more intensively studied. Our results, which are based on the largest number of bird and mammal species compiled to date, bring convincing support that the process of reproductive senescence is embedded in the life-history strategy of a given species [5,20,21] and has a role in the evolution of life histories.

The degree of sociality appears to have a very limited direct influence on reproductive senescence when the effects of allometry and pace of life are accounted for, which supports the view expressed above that reproductive senescence in a given species is mostly driven by the species size and position on the slow-fast continuum of life histories. With the exception of cooperative breeding in mammals, none of the sociality traits we analysed (i.e. relative brain size in birds and mammals, colonial breeding and parental cooperation in birds) were associated with either the occurrence or the rate and onset of reproductive senescence. These results support the conclusions reached about the putative role of sociality in the evolution of actuarial senescence and longevity [7].

One striking result of this work is that the degree of sociality was associated with a decreased strength of senescence in terms of both rate and onset when species differences in pace of life were not controlled for. As these associations vanished when we controlled for the pace of life, we conclude that the social mode of life *per se* does not influence reproductive senescence. Instead, the social lifestyle seems to shape the entire life-history strategy, which supports H2 and refutes H1. Cooperative breeders often display delayed dispersal and reproductive suppression of subordinates [45], so that the age at first reproduction is also delayed and the number of breeding attempts is thus decreased, which can lead to increased longevity [20]. Moreover, evidence suggests that a slower pace of life is evolutionary linked to colonial breeding in birds [40] and to larger brain size in mammals [66], and species displaying a high degree of sociality also display slower development, delayed age at primiparity, better survival prospects and longer lifespan (reviewed in [7]). Whether a large relative brain size is directly related to a slower pace of life (cognitive buffer hypothesis; [67]) for a given degree of sociality or a large relative brain size is more likely to evolve in social species (social brain hypothesis; [41]), which leads to slow down the pace of life is currently unknown and requires further investigation. However, we cannot rule out the alternative hypothesis (H3) that pace of life has independent effects on both social lifestyle and reproductive senescence, involving the absence of a functional link between reproductive senescence and the degree of sociality. The current view is that sociality shapes the evolution of life histories and senescence [7]. However, consistent with the alternative hypothesis, there is evidence in birds showing that species in which a slow pace of life have evolved (i.e. long life) are more prone to evolve a social lifestyle (cooperative breeding) [68]. Therefore, from our findings, we can likely reject a direct association between reproductive senescence and degree of sociality (H1), but whether they are indirectly related through the shaping of senescence by the pace of life (H2) or simply independent responses to the pace of life (H3) cannot be assessed. Future studies using a phylogenetic path analysis or ancestral character reconstruction approach for sociality, life history and senescence traits could differentiate between the latter two alternatives. This analysis will require much improved metrics of the degree of sociality.

It might be premature to conclude firmly that the degree of sociality has no direct effect on the magnitude of reproductive senescence. Currently, we lack accurate metrics for measuring the degree of sociality across a wide range of species and the metrics we used in this study have limitations. For instance, cooperative breeding would require a more detailed typology based on four classes (i.e. solitary, social, communal and cooperative; as per [51]) to describe accurately the different levels of social complexity. Moreover, because all colonial species in our data set are seabirds and occupy thus marine (aquatic) habitats, colonial breeding might be confounded by habitat type if aquatic species evolve slower pace of life irrespective of coloniality. However, contrary to this expectation, terrestrial organisms generally have a slower pace of life than aquatic ones [69], which suggests that coloniality might play a role in the evolution of pace of life without being confounded by habitat type. Nevertheless, future studies will be required to assess whether the association between coloniality and reproductive senescence differs (with and without accounting for the pace of life) between terrestrial and marine colonial species. Unfortunately, we failed to identify any data fulfilling our selection criteria on reproductive senescence in terrestrial colonial birds. We also relied on the social brain hypothesis, which proposes that relative brain size is larger in species with a higher degree of social bonds [41], to justify our use of the relative brain size as a measure of the degree of sociality. This hypothesis has received so far mixed support when assessing its plausibility in animal taxa with a wide diversity of social systems [41,70]. However, the social brain hypothesis holds for species displaying complex social interactions, such as cetaceans or primates [41,71]. Our results based on relative brain size should also be treated with caution because brain size is only a rough index of sociality and is related to other life-history traits that might influence senescence (e.g. relationship with longevity [72,73]). Taken together, our conclusion that the degree of sociality has no direct influence on reproductive senescence in birds and mammals will need to be investigated more thoroughly when better measures of the degree of sociality will be available for a substantial set of species. The recent development of social network analysis [74], which allows detailed accounts of individual interactions within populations, should play a key role for doing that.

Interestingly, the only detectable direct effect of the degree of sociality on reproductive senescence was opposite to our prediction. We found that cooperatively breeding mammals senesce faster, not slower, than non-cooperative ones for a given size and pace of life. At first sight, this finding contradicts within-population studies that showed almost consistently that helpers buffer the demographic senescence of breeders [7,45]. However, cooperative breeding might have opposite effect on reproductive senescence depending on the level of biological organization we consider. For instance, getting the breeder tenure requires winning aggressive social interactions that increase the level of physiological stress at the long term [75,76], which might exacerbate reproductive senescence [8]. Additionally, the buffered effect of cooperative breeding on reproductive senescence among individuals within a population of a given species can translate into an increased reproductive senescence of cooperative breeding species compared with non-cooperative breeding ones. Within a population of cooperative breeders, reproductively active individuals (i.e. dominants) usually receive alloparental assistance from helpers (i.e. subordinates), which decreases the cost of a given reproductive effort and leads thereby either to a postponed onset or to a decelerated rate of reproductive or actuarial senescence (e.g. load-lightening hypothesis; [24]). For instance, in Alpine marmots (*Marmota marmota*), individuals that have benefited from more helping during their prime-age reproductive stage display a reduced actuarial senescence compared to those that received less help [77], leading to increased individual heterogeneity in the amount of senescence. At the population level, as considered in across-species analyses, the reproductive suppression and associated physiological stress of individuals that help repeatedly before reaching a dominant status and/or the paucity of substantial help when being breeders might lead to more pronounced reproductive senescence. Overall, at the population level, increased costs of helping when subordinate or lack of help when dominant for a large number of individuals might counterbalance the benefits of having many helpers during breeding events that only a reduced number of individuals enjoy. This strong individual heterogeneity in the strength of reproductive senescence within populations of cooperative breeders might lead the average magnitude of reproductive senescence to be higher in these species than in non-cooperatively breeding ones. An alternative explanation is that females of cooperatively breeding mammals have higher reproductive output, which, for a given pace of life, ultimately results in higher rate of reproductive senescence. Indeed, in mammal species in which females receive offspring provisioning help from males, females have higher reproductive output (larger litter size and shorter inter-birth intervals; [78,79]).

## 5. Conclusions

Our results indicate that degree of sociality is not directly associated with female reproductive senescence. Instead, the positive covariation between the degree of sociality and a slower pace of life has deeper evolutionary roots, which encompass both a later onset and a slower rate of reproductive senescence.

## Ethics

This study does not have ethical aspects.

## Data accessibility

The data sets supporting this article have been uploaded as part of the Electronic Supplementary Material (see ESM ‘Data set’).

## Authors’ contributions

C.I.V., J.-F.L. and J.-M.G. designed the study; C.I.V., J.-F.L., O.V., P.L.P., V.R. and J.-M.G. provided data; O.V. and V.R. carried out the statistical analyses; C.I.V., O.V., J.-F.L. and J.-M.G. drafted the manuscript with considerable help from P.L.P. and V.R.; all authors approved the final version of the manuscript.

## Acknowledgements

We are grateful to Hanna Kokko and three anonymous reviewers for insightful comments that greatly improved our manuscript.

