## Additional details for Methods and Results for "Is degree of sociality associated with reproductive senescence? A comparative analysis across birds and mammals" (from another version of this preprint: version 1, 2020-10-30)

ELECTRONIC SUPPLEMENTARY MATERIAL – ESM

Running title: Sociality and reproductive senescence

ORCID: 0000-0002-8736-2391 (CIV), 0000-0001-5789-2124 (OV), 0000-0001-9898-2353 (JFL), 0000-0002-3659-7684 (PLP), 0000-0001-9991-0482 (VR), 0000-0003-0174-8451 (JMG)

2. Methods

(a) Reproductive senescence data

We extracted data similar to those obtained in mammals [1] by the systematic search of the literature reporting reproductive senescence data in birds. We used the following keyword strings and boolean operator for literature search:

(“reproductive senescence” OR “reproductive aging” OR “reproductive ageing” OR **“age-related reproductive success” OR “age-specific reproductive success”** OR **“age-related fertility” OR “age-specific fertility” OR “age-related fecundity” OR “age-specific fecundity”** OR **“age-related hatching success” OR “age-specific hatching success” OR “age-related fledging success” OR “age-specific fledging success”** OR **“age-related reproductive effort” OR “age-specific reproductive effort”** OR **“age-related reproductive output” OR “age-specific reproductive output”)**,

both in Scopus and Web of Science. Scopus provided 2,480 hints (access date: 6th May, 2020), while Web of Science yielded 2,042 hints (access date: 14th May, 2020) of which 1,380 hints were identified as duplicates between the two sources. We also surveyed the scientific publications citing two seminal papers dealing with the evolution of ageing [2,3] and a book about the evolution of life histories in birds [4]. Finally, we did additional search for data in Google Scholar, ecological and ornithological journals, and by snow-balling.

(b) Life-history and sociality traits

**Table S1.** Pair-wise correlation between life-history traits that were used to extract a size component and a pace of life component (see phylogenetic principal component analyses in the main text) and brain size in both birds and mammals. The correlation coefficient is provided with the sample size in parentheses. Each variable is log-transformed. FBM – female body mass, AFR – age at first reproduction, LON – longevity, BRM – brain mass.

|  | birds | | |  | Mammals | | |
| --- | --- | --- | --- | --- | --- | --- | --- |
|  | log AFR | log LON | log BRM |  | log AFR | log LON | log BRM |
| log FBM | 0.66 (36) | 0.76 (36) | 0.98 (22) |  | 0.63 (101) | 0.68 (101) | 0.97 (88) |
| log AFR |  | 0.84 (36) | 0.67 (22) |  |  | 0.88 (101) | 0.72 (88) |
| log LON |  |  | 0.85 (22) |  |  |  | 0.79 (88) |

We analysed female body mass, age at first reproduction and longevity using a phylogenetically-controlled principal component analysis (PPCA) to extract a size component (i.e. first PPCA axis, hereafter PPC size) and a pace of life component (i.e. second PPCA axis, hereafter PPC pace). PPC size is defined by the positive covariation among body mass, age at first reproduction and longevity and was used to control for allometry, while PPC pace is defined by the positive covariation between age at first reproduction and longevity for a given size and was used to control for pace of life.

**Table S2.** Summary of the two PPCA analyses performed separately for birds and mammals. Variance components of each principal component are given along with loads of individual variables included in the PPCA. PPC1 is a size component (hereafter PPC size) and PPC2 is a pace of life component (hereafter PPC pace).

|  |  | birds | | | mammals | | |
| --- | --- | --- | --- | --- | --- | --- | --- |
|  |  | PPC1 | PPC2 | PPC3 | PPC1 | PPC2 | PPC3 |
| variance | standard deviation | 0.19 | 0.11 | 0.07 | 0.28 | 0.12 | 0.09 |
|  | proportion of variance | 0.69 | 0.23 | 0.09 | 0.79 | 0.12 | 0.08 |
|  | cumulative proportion | 0.69 | 0.91 | 1.00 | 0.79 | 0.92 | 1.00 |
| loads | log female body mass | 0.97 | 0.26 | –0.04 | 1.00 | 0.08 | –0.02 |
|  | log age at first reproduction | 0.53 | –0.80 | –0.29 | 0.40 | –0.90 | –0.14 |
|  | log longevity | 0.61 | –0.48 | 0.63 | 0.34 | –0.23 | 0.91 |

3. Results

**Table S3.** Models of occurrence, rate and onset of reproductive senescence in birds. Indented models show base models to which all models beneath are compared using Akaike Information Criterion (AIC). To illustrate the effect of explanatory variables in the base models, the effect of removing these from the model are also presented. For models beneath the base model, only the removal (–) or addition (+) of explanatory terms are shown. Models on the left show models with pace of life included, while those on the right show models without pace of life. The effect of polynomial predictors is presented by replacing polynomial terms with a linear one or vice versa. Note that pace of life was not a statistically significant predictor of occurrence of reproductive senescence; therefore, this model was not repeated without including pace of life as an explanatory variable. Only models based on the same sample size are comparable. Note that ΔAIC < |2| are comparable in fit and in this case we preferred the most parsimonious model. Social traits are italicized and those with statistically significant effect are italicized and marked in bold.

| including pace of life (PPC pace) | | | |  | excluding pace of life (PPC pace) | | | |
| --- | --- | --- | --- | --- | --- | --- | --- | --- |
| base model | *n* | AIC | ΔAIC |  | base model | *n* | AIC | ΔAIC |
| occurrence of senescence ~ 1 | 36 | 49.04 | 0.00 |  |  |  |  |  |
| ~ + PPC size | 36 | 48.85 | –0.19 |  |  |  |  |  |
| ~ + poly(PPC size, 2) | 36 | 49.50 | 0.47 |  |  |  |  |  |
| ~ + PPC pace | 36 | 51.04 | 2.01 |  |  |  |  |  |
| ~ + poly(PPC pace, 2) | 36 | 52.56 | 3.52 |  |  |  |  |  |
| ~ + reproductive trait | 36 | 48.56 | –0.47 |  |  |  |  |  |
| *~ + coloniality* | 36 | 50.62 | 1.58 |  |  |  |  |  |
| occurrence of senescence ~ 1 | 22 | 33.35 | 0.00 |  |  |  |  |  |
| *~ + residual brain size* | 22 | 33.05 | –0.31 |  |  |  |  |  |
| occurrence of senescence ~ 1 | 31 | 44.72 | 0.00 |  |  |  |  |  |
| *+ parental cooperation* | 31 | 46.73 | 2.01 |  |  |  |  |  |
| base model | *n* | AIC | ΔAIC |  | base model | *n* | AIC | ΔAIC |
| log rate of senescence ~ poly(PPC size, 2) + PPC pace + reproductive trait | 22 | 53.71 | 0.00 |  | log rate of senescence ~ poly(PPC size, 2) + reproductive trait | 22 | 65.05 | 0.00 |
| ~ – poly(PPC size,2) | 22 | 65.52 | 11.81 |  | ~ – poly(PPC size,2) | 22 | 69.02 | 3.97 |
| ~ – poly(PPC size,2) + PPC size | 22 | 56.79 | 3.08 |  | ~ – poly(PPC size,2) + PPC size | 22 | 68.76 | 3.71 |
| ~ – reproductive trait | 22 | 58.11 | 4.40 |  | ~ – reproductive trait | 22 | 67.14 | 2.09 |
| ~ – PPC pace | 22 | 65.05 | 11.34 |  |  |  |  |  |
| ~ – PPC pace + poly(PPC pace,2) | 22 | 52.74 | –0.97 |  |  |  |  |  |
| ~ + log onset of senescence | 22 | 55.71 | 2.00 |  | ~ + log onset of senescence | 22 | 63.64 | –1.41 |
| ~ + standardized (yes) | 22 | 55.14 | 1.43 |  | ~ + standardized (yes) | 22 | 65.76 | 0.70 |
| *~ + coloniality* | 22 | 55.57 | 1.86 |  | ***~ + coloniality*** | **22** | **57.93** | **–7.13** |
| log rate of senescence ~ poly(PPC size, 2) + PPC pace + reproductive trait | 12 | 32.27 | 0.00 |  | log rate of senescence ~ poly(PPC size, 2) + reproductive trait | 12 | 32.91 |  |
| *~ + residual brain size* | 12 | 34.19 | 1.92 |  | *~ + residual brain size* | 12 | 34.19 | 1.28 |
| log rate of senescence ~ poly(PPC size, 2) + PPC pace + reproductive trait | 18 | 49.95 | 0.00 |  | log rate of senescence ~ poly(PPC size, 2) + reproductive trait | 18 | 49.95 | 0.00 |
| *~ + parental cooperation* | 18 | 51.58 | 1.63 |  | *~ + parental cooperation* | 18 | 51.58 | 1.64 |
| base model | *n* | AIC | ΔAIC |  | base model | *n* | AIC | ΔAIC |
| log onset of senescence ~ PPC size + PPC pace | 22 | 30.19 | 0.00 |  | log onset of senescence ~ PPC size | 22 | 36.15 | 0.00 |
| ~ – PPC size | 22 | 45.19 | 15.00 |  | ~ – PPC size | 22 | 44.44 | 8.29 |
| ~ – PPC size + poly(PPC size,2) | 22 | 34.09 | 3.90 |  | ~ – PPC size + poly(PPC size,2) | 22 | 37.31 | 1.16 |
| ~ – PPC pace | 22 | 36.15 | 5.96 |  |  |  |  |  |
| ~ – PPC pace + poly(PPC pace, 2) | 22 | 28.57 | –1.62 |  |  |  |  |  |
| ~ + reproductive trait | 22 | 28.67 | –1.52 |  | ~ + reproductive trait | 22 | 39.43 | 3.28 |
| *~ + coloniality* | 22 | 32.17 | 1.99 |  | ***~ + coloniality*** | **22** | **34.10** | **–2.05** |
| log onset of senescence ~ PPC size + PPC pace | 12 | 9.95 | 0.00 |  | log onset of senescence ~ PPC size | 12 | 17.52 | 0.00 |
| *~ + residual brain size* | 12 | 11.53 | 1.58 |  | *~ + residual brain size* | 12 | 17.93 | 0.41 |
| log onset of senescence ~ PPC size + PPC pace | 18 | 28.21 | 0.00 |  | log onset of senescence ~ PPC size | 18 | 30.56 | 0.00 |
| *~ + parental cooperation* | 18 | 29.39 | 1.18 |  | *~ + parental cooperation* | 18 | 29.88 | –0.68 |

**Table S4.** Models of occurrence, rate and onset of reproductive senescence in mammals. Only full models are presented. Models on the right show models with pace of life removed. Social traits are italicized and those with statistically significant effect are italicized and marked in bold.

|  | including pace of life (PPC pace) | | | |  | excluding pace of life (PPC pace) | | | | |
| --- | --- | --- | --- | --- | --- | --- | --- | --- | --- | --- |
| occurrence of senescence |  | *β* (s.e.) | *z* | *p* |  |  | *β* (s.e.) | | *z* | *p* |
|  | intercept | 1 (0.44) | 2.26 | 0.0236 |  | Intercept | 1.13 (0.43) | | 2.62 | 0.0088 |
|  | quality (transversal) | –1.29 (0.51) | 2.51 | 0.0119 |  | quality (transversal) | –1.29 (0.52) | | 2.48 | 0.0132 |
|  | hunted (yes) | –0.31 (0.6) | 0.51 | 0.6074 |  | hunted (yes) | –0.4 (0.6) | | 0.67 | 0.5058 |
|  | PPC size | 0.23 (0.1) | 2.18 | 0.0294 |  | PPC size | 0.23 (0.1) | | 2.27 | 0.0233 |
|  | PPC pace | –0.01 (0.4) | 0.03 | 0.9755 |  |  |  | |  |  |
|  | *residual brain size* | 1.01 (0.64) | 1.59 | 0.1129 |  | *residual brain size* | 0.99 (0.61) | | 1.62 | 0.1062 |
|  | *cooperative breeding (yes)* | 0.1 (0.82) | 0.12 | 0.9077 |  | *cooperative breeding (yes)* | 0.02 (0.83) | | 0.02 | 0.9855 |
|  | model stats: *α* = 0.0434, AIC = 109.46, *n* = 88 | | | |  | model stats: *α* = 0.0544, AIC = 107.28, *n* = 88 | | | | |
| log rate of senescence |  | *β* (s.e.) | *t* | *P* |  |  | *β* (s.e.) | | *t* | *p* |
|  | intercept | –2.12 (0.59) | 3.58 | 0.0008 |  | intercept | –1.6 (0.54) | | 2.97 | 0.0045 |
|  | log onset of senescence | –0.03 (0.27) | 0.12 | 0.9083 |  | log onset of senescence | –0.33 (0.24) | | 1.37 | 0.1758 |
|  | quality (transversal) | –0.06 (0.28) | 0.21 | 0.8370 |  | quality (transversal) | –0.07 (0.3) | | 0.23 | 0.8170 |
|  | hunted (yes) | 0.54 (0.35) | 1.56 | 0.1260 |  | hunted (yes) | 0.56 (0.36) | | 1.57 | 0.1220 |
|  | PPC size | –0.43 (0.06) | 6.73 | < 0.0001 |  | PPC size | –0.44 (0.08) | | 5.37 | < 0.0001 |
|  | poly(PPC pace, 2)1 | –4.45 (1.35) | 3.28 | 0.0019 |  |  |  | |  |  |
|  | poly(PPC pace, 2)2 | –2.41 (1.08) | 2.24 | 0.0295 |  |  |  | |  |  |
|  | *residual brain size* | –0.14 (0.31) | 0.46 | 0.6483 |  | ***residual brain size*** | **–0.64 (0.32)** | | **2.03** | **0.0475** |
|  | ***cooperative breeding (yes)*** | **1.3 (0.43)** | **3.05** | **0.0037** |  | *cooperative breeding (yes)* | 0.85 (0.49) | | 1.71 | 0.0927 |
|  | model stats: *λ* = 0.0417, AIC = 161.09, *n* = 58 | | | |  | model stats: *λ* = 0.2794, AIC = 171.72, *n* = 58 | | | | |
| log onset of senescence |  | *β* (s.e.) | *t* | *p* |  |  | *β* (s.e.) | | *t* | *p* |
|  | intercept | 1.81 (0.18) | 10.33 | **< 0.0001** |  | intercept | 1.87 (0.24) | | 7.63 | < 0.0001 |
|  | quality (transversal) | –0.17 (0.13) | 1.3 | 0.1995 |  | quality (transversal) | –0.17 (0.14) | | 1.18 | 0.2444 |
|  | hunted (yes) | 0.15 (0.16) | 0.94 | 0.3494 |  | hunted (yes) | 0.04 (0.17) | | 0.21 | 0.8311 |
|  | PPC size | 0.18 (0.04) | 4.8 | < 0.0001 |  | PPC size | 0.23 (0.05) | | 5 | < 0.0001 |
|  | PPC pace | 0.54 (0.13) | 4.11 | 0.0001 |  |  |  | |  |  |
|  | *residual brain size* | 0.1 (0.17) | 0.56 | 0.5811 |  | ***residual brain size*** | **0.38 (0.19)** | | **2.01** | **0.0499** |
|  | *cooperative breeding (yes)* | –0.13 (0.25) | 0.55 | 0.5861 |  | *cooperative breeding (yes)* | 0.12 (0.26) | | 0.47 | 0.6375 |
|  | model stats: *λ* = 0.6053, AIC = 85.94, *n* = 58 | | | |  | model stats: *λ* = 0.8102, AIC = 98.5, *n* = 58 | | | | |
